# Microparticle-based Biochemical Sensing Using Optical Coherence Tomography and Deep Learning

**DOI:** 10.1101/2020.12.21.422771

**Authors:** Shreyas Shah, Chun-Nam Yu, Mingde Zheng, Heejong Kim, Michael S. Eggleston

## Abstract

Advancing continuous health monitoring beyond vital signs to biochemistry will revolutionize personalized medicine. Herein, we report a novel platform to achieve remote biochemical monitoring using microparticle-based biosensors and optical coherence tomography (OCT). Stimuli-responsive, polymeric microparticles were designed to serve as freely-dispersible biorecognition units, wherein binding with a target biochemical induces volumetric changes of the microparticle. Analytical approaches to detect these sub-micron changes in 3D using OCT were devised by modeling the microparticle as an optical cavity, enabling estimations far below the resolution of the OCT system. As a proof of concept, we demonstrated the 3D spatiotemporal monitoring of glucose-responsive microparticles distributed throughout a tissue-mimic in response to dynamically-fluctuating levels of glucose. Deep learning was further implemented using 3D convolutional neural networks to automate the vast processing of the continuous stream of three-dimensional time series data, resulting in a robust end-to-end pipeline with immense potential for continuous *in vivo* biochemical monitoring.

## INTRODUCTION

Biological signals emanating from the human body are important indicators for tracking overall health and well-being. Developing innovative methods to measure these diverse signals has played an essential role in pushing the frontiers of healthcare. Due to advances in wearable technologies, measuring vital signs (e.g. heart rate, body temperature, respiration rate, blood pressure) is now commonplace in the comforts of our very homes^1^. However, tapping into the rich in-body biochemistry for continuous and real-time monitoring has proven to be much more challenging. Acquiring a continuous readout at the biomolecular level can provide a much more informed assessment of an individual’s state of well-being^2^. Although continuous glucose monitoring is well-established and commercially available^3^, detecting and monitoring other types of biomarkers in an *in vivo* transdermal setting has remained elusive^4^. This is partly due to challenges in developing strategies which can couple the complex binding activity of a target biomarker with a sensing modality to spatiotemporally monitor the binding within tissue over time^5^.

In terms of a robust sensing modality, optical methods have proven to be promising for continuous physiological monitoring. Compared to other types of sensors (e.g. electrical, acoustic), optical-based biosensors offer several advantages including high sensitivity, wide dynamic range for detection, immunity from electromagnetic interference, capability for multiplexing with different wavelengths of light, and non-contact means for interrogation^6^. Conventional optical readouts tend to be generated as a colorimetric, fluorescent or luminescent response; fluorescence-based methods are most common for *in vivo* biosensing^7-10^. Although a number of studies have reported the use of organic dyes to detect fluorescence modulation in response to biochemical binding^11-14^, there are several limitations for fluorescence-based biosensing in the context of long-term monitoring, which include: susceptibility to photobleaching, low signal-to-noise intensity due to tissue autofluorescence, low penetration depths for visible light excitation, and a limited chemical versatility in fluorescent probe design to sense diverse biochemical targets^15^.

Unlike these conventional optical-based readouts, optical coherence tomography (OCT) can offer significant advantages for continuous *in vivo* biosensing. By utilizing infrared light, OCT enables depth-resolved cross-sectional imaging of tissue in a non-contact and non-invasive manner^16^. Moreover, detailed microscopic structures can be resolved in three dimensions with a high spatial resolution of ~5-10 μm^17^. OCT also operates at much faster speeds than other tomography imaging techniques (like ultrasound) since it is based on light, providing 3D imaging at a temporal resolution of about one second. Beyond acquiring structural information, depth-resolved spectroscopic information can also be extracted from the OCT signal using short-time Fourier transforms, further providing invaluable biochemical insight about endogenous biomolecules native to the tissue or exogenous agents introduced into the tissue^18,19^.

Previous studies have attempted to exploit these features of OCT for biosensing. Since glucose can alter the optical properties of tissues, OCT has been used to correlate changes in tissue scattering coefficients to blood glucose concentrations^20,21^. However, this approach has proven to be unreliable and inaccurate, since the tissue scattering coefficients can also vary due to subtle changes in body temperature and the concentration of other analytes. To enhance specificity, several studies designed biomolecular recognition devices to monitor changes in turbidity in response to changing biochemical concentration^22,23^. However, such approaches faced challenges with implanting large devices (e.g. gold mirror surfaces) and precisely positioning the device in the tissue with respect to the laser beam. Spectroscopic OCT has also been employed to characterize the optical spectra of micron-scale particles within a tissue^24,25^. However, this approach only worked to estimate nuclear sizes of static targets and labels within the native tissue rather than as a dynamic biosensing modality.

Herein, we report a novel biosensing approach that combines the key attributes inherent to OCT for the dynamic monitoring of biochemical-responsive, tissue-embeddable microparticles (**Fig. 1**). Polymeric microparticles were engineered to serve as freely dispersible biomolecular recognition units, wherein changes in the binding of a biomolecule would induce volumetric changes of the microparticle. By employing glucose-responsive microparticle biosensors as a proof of concept, we demonstrate 3D spatial tracking of the micron-scale biosensors distributed throughout a hydrogel-based tissue-mimic as well as temporal monitoring of the physical and spectral changes of the distributed biosensors in response to a dynamically-fluctuating biochemical microenvironment. The physical and spectral changes in the microparticle size were monitored using two distinct analytical approaches, which enabled estimations far below the resolution of the OCT system. In order to overcome the significant user input required to manually locate the biosensor response, we further developed a deep learning-based approach to automate the vast processing of the continuous stream of three-dimensional time series data. Overall, we demonstrate an end-to-end OCT-based biosensing pipeline, which has immense potential for realizing continuous and real-time *in vivo* monitoring of physiologically-relevant biomarkers.

**Figure 1.**
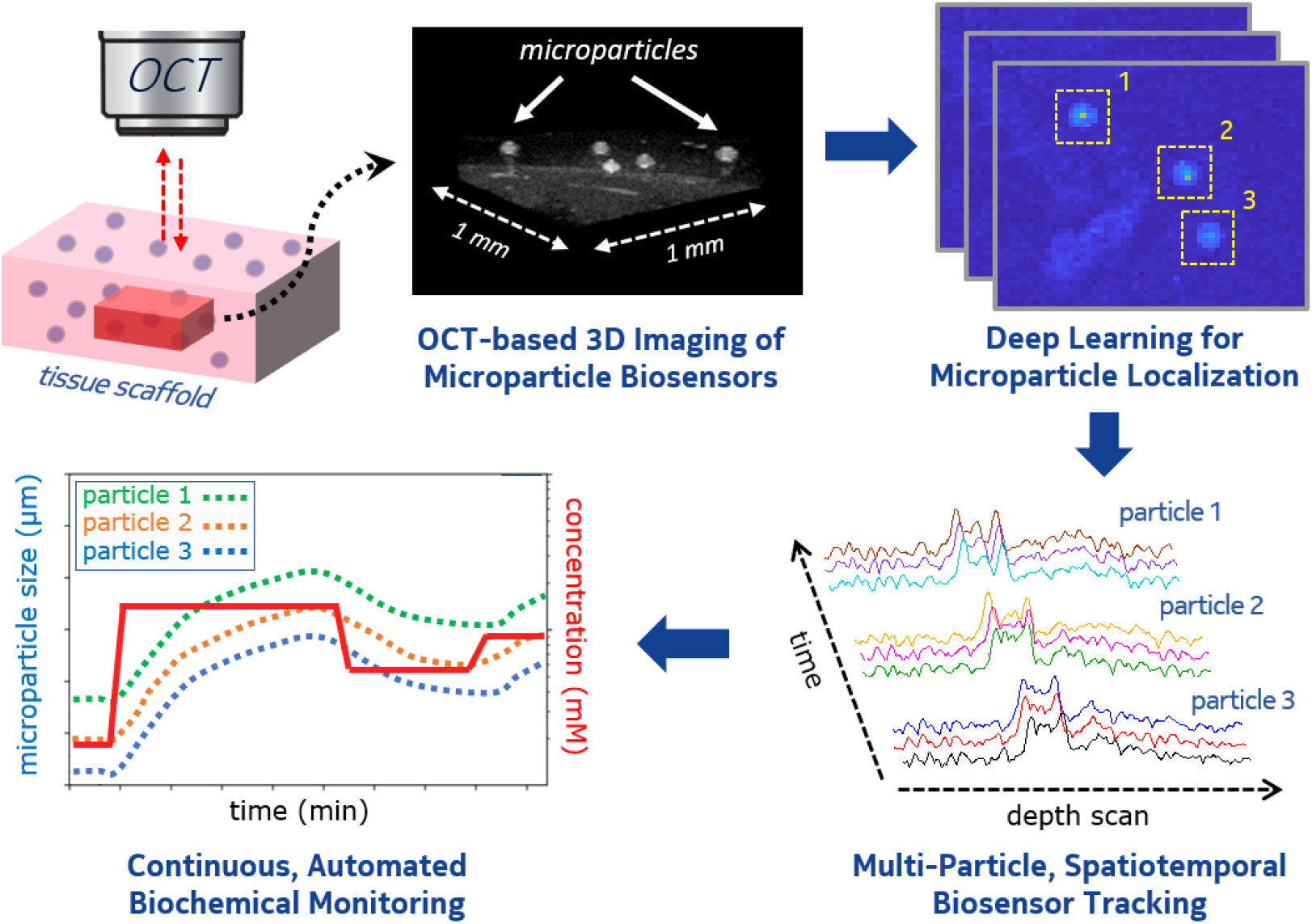
Schematic diagram depicting the end-to-end pipeline for OCT-based automated biochemical monitoring using embeddable, biochemical-responsive microparticle biosensors.

## RESULTS & DISCUSSION

### Designing hydrogel-based spherical microparticles

Stimuli-responsive hydrogels are known to alter their physical and chemical properties upon exposure to an external stimulus like light, temperature, enzymes and biochemicals^26^. These responsive biomaterials have been extensively utilized over the years for numerous applications including tissue engineering, drug delivery and biosensing^27^. We sought to fabricate glucose-responsive microparticles by covalently incorporating a phenylboronic acid derivative into a hydrogel matrix. Previous studies have demonstrated glucose-responsive hydrogels in the form of films and fibers, wherein the reversible complexation of the *cis*-diol groups of glucose molecules with the phenylboronic acid derivative was shown to increase the fraction of charged boronate species, which in turn increased the osmotic pressure within the hydrogel and induced volumetric swelling^28,29^. In order to achieve isotropic swelling and deswelling synchronized to variations in glucose concentration, we adopted a similar sensing mechanism to optimize the generation of spherical hydrogel microparticles (**Fig. 2a**). We incorporated the 3-(acrylamido)phenylboronic acid (APBA) molecule as the glucose-responsive element into a copolymer backbone composed of poly(acrylamide-*co*-poly(ethylene glycol) diacrylate) p(AM-*co*-PEGDA) (**Supplementary Fig. 1**).

**Figure 2.**
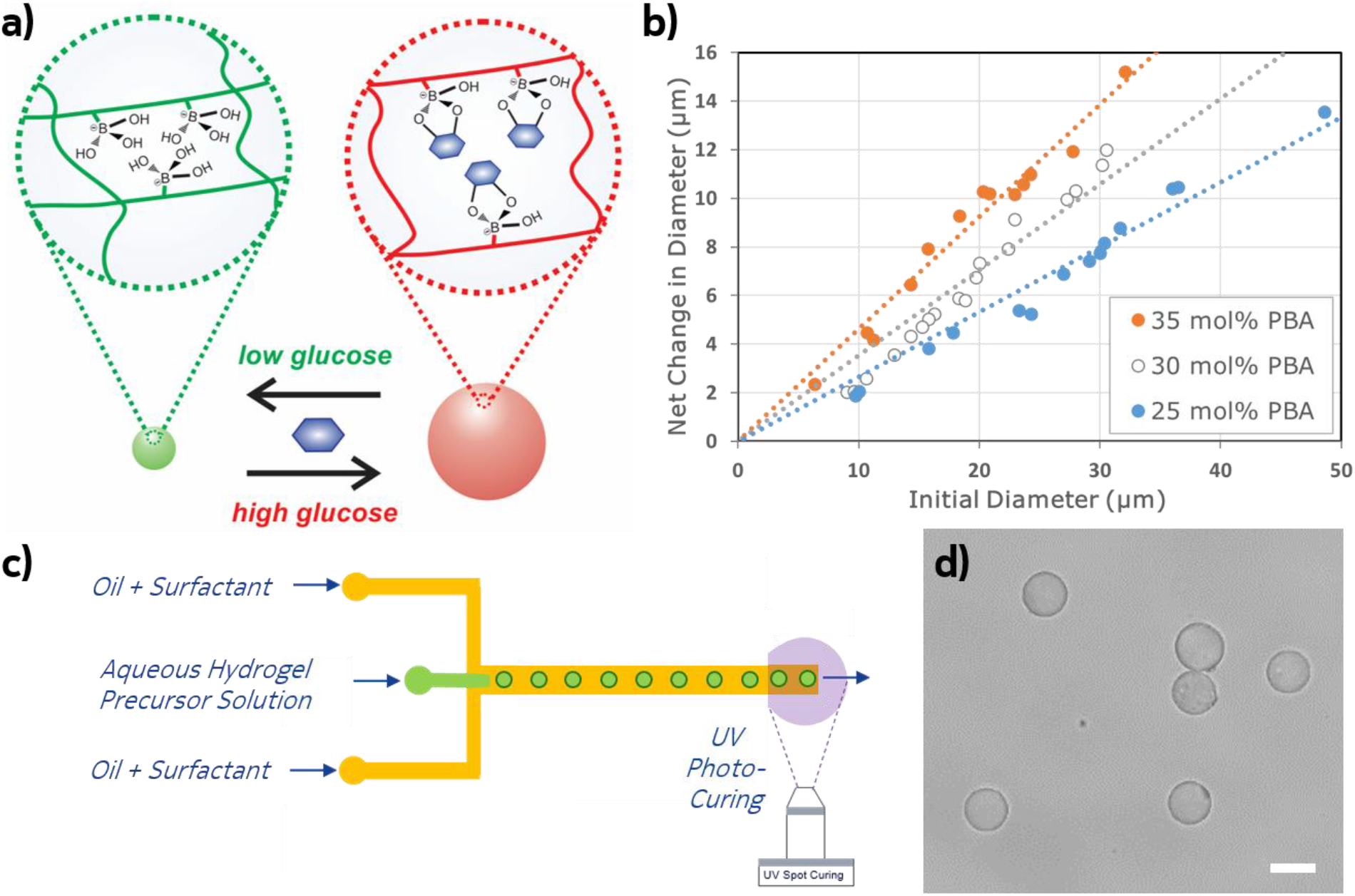
Synthesis and characterization of biochemical-responsive hydrogel microparticles. (a) The hydrogel microparticle consists of a PEG-crosslinked polyacrylamide backbone functionalized with 3-acrylamidophenylboronic acid. The phenylboronic acid derivative binds the cis diols of glucose molecules, leading to a reversible binding reaction that modulates the osmotic pressure and results in volumetric changes in the microparticle size. (b) Plot depicting the net change in microparticle size versus the initial microparticle diameter for varying doping concentrations of 3-(acrylamide)phenylboronic acid. (c) Droplet microfluidic setup for microparticle synthesis, achieved through co-flow of the aqueous hydrogel precursor solution and the oil phase followed by UV-based photo-crosslinking. (d) Microscopy images of microfluidic-generated monodisperse microparticles. Scale bar: 25μm.

Microparticles were generated using an emulsification process, wherein droplets containing the monomers (i.e. AM, PEGDA and APBA) in an aqueous medium were dispersed in an immiscible medium (i.e. oil). Stable emulsion droplets favor a spherical shape due to the minimized interfacial energy at the boundary between the two immiscible fluids and the retention of uniform pressure over the entire interface^30^. The emulsion droplets can then be polymerized to form spherical hydrogel microparticles; in our case, we added a photoinitiator (2-hydroxy-2-methylpropiophenone) to the initial precursor monomer solution, rendering the droplets photo-crosslinkable upon exposure to UV light (365 nm).

### Fabrication and testing of glucose-responsive microparticles

Emulsions were formed using one of two distinct synthetic approaches. The first method involved vortexing, in which the aqueous monomer solution was deposited in oil and then submitted to bulk shearing forces. Although the average microparticle size range can be roughly controlled by adjusting the vortexing speed, the size distribution is relatively broad (**Supplementary Fig. 2**). Nevertheless, the vortexing method proved to be advantageous for investigating the glucose-sensing properties of different-sized microparticles for a given chemical composition (**Fig. 2b**). In general, larger microparticles would tend to have a larger amount of APBA available for binding, leading to a greater increase in microparticle size in response to glucose. This is observed in **Fig. 2b**, in which the net change in the microparticle size upon exposure to glucose showed a linear relationship with the initial starting size of the microparticle. However, by tuning the relative concentration of APBA and AM, varying degrees of maximal size change can be achieved. As the concentration of APBA increased from 25mol% to 35mol%, there was a corresponding increase in the net size change at each given initial starting size (**Fig. 2b**). While increasing the concentration of APBA can achieve a greater response, it is counterbalanced by requiring more time to reach equilibrium. For instance, upon exposure to 100mM of glucose, an expansion in microparticle size of ~25% was achieved within three minutes with a composition of 25mol% APBA, ~32% expansion within five minutes at 30mol% APBA, and ~45% expansion within eight minutes at 35mol% APBA (**Supplementary Table 1**). Considering the maximum glucose response and the time required to reach equilibrium, we found 30mol% to be the optimum concentration of APBA, and used the composition 67:3:30 of AM:PEGDA:APBA for the subsequent experiments.

After optimizing the desired chemical composition of the microparticle, we employed microfluidic flow-focusing as a second synthetic method to prepare monodisperse emulsions that yield highly uniform microparticles of a desired size (**Fig. 2c**). Microfluidic flow-focusing is a popular and versatile technique^31^, in which the desired droplet size can be achieved by adjusting the dimension of the orifice, as well as tuning the flow rates of the dispersal and continuous phase fluids. By designing a microfluidic channel orifice having 10μm or less in width and depth, hydrodynamically dispersed droplets can achieve sizes larger than 20μm at a fast rate with high yield. Moreover, because fluidic stability is a key to sustained microfluidic generation with uniform sizes, this process can be optimized by using an isolated chamber to minimize sources of potential flow interruptions such as mechanical interferences from tubing drag and stage movement. Droplets of the hydrogel precursor were formed as an assembly line, achieved through co-flow of the aqueous hydrogel precursor phase and an oil phase containing surfactants (**Supplementary Fig. 3**). We adjusted the flow rate (0.25 - 40μL/min) to acquire distinct size populations between 20 to 60μm. The delivery of UV light for the photo-polymerization of the droplet emulsions was achieved through a high-power microscope lens array and built-in UV source. This setup permitted targeted spot-curing of the droplets during transit through the microfluidic device, thereby ensuring the desired microparticle shape and size. The microfluidic emulsification approach allowed for the synthesis of uniform, monodisperse microparticles (**Fig. 2d**). The glucose response of microparticles synthesized using this approach was further confirmed at physiologically-relevant glucose concentrations (**Supplementary Fig. 4**).

### Flow cell setup to mimic the dynamic *in vivo* microenvironment

Microparticles were embedded in a tissue-mimic composed of agarose, and a thin layer was deposited in a polydimethylsiloxane (PDMS)-based flow cell device. The device was then placed in a fixed position under the scan lens (**Supplementary Fig. 5**). The impinging jet of the analyte (i.e. glucose) was delivered using syringe pumps in a flow parallel to the embedded microparticle sensors, wherein advection and diffusion allowed for the transport of glucose throughout the flow cell. This experimental setup allowed us to monitor the microparticle response under a continuous flow of fluctuating levels of glucose. We initially monitored the microparticles over time using optical microscopy. The microparticles were observed to undergo swelling when the glucose concentration increased and shrinkage when the glucose concentration decreased (**Fig. 3a**). In this case, the microparticle size was calculated at each time point using manual segmentation and estimation of the microparticle diameter in 2D images. The subtle variations over time in microparticle size were detectable on a continuous basis within the normal physiological range of glucose (4-6mM) as well as levels indicative of hyperglycemia (>10mM)^32^ (**Fig. 3b**). In this way, the microparticles serve as an adequate proof of concept biosensor which can be freely distributed within a 3D tissue-like matrix. However, optical microscopy is limited to 2D imaging of thin surfaces and optically transparent mediums. It offers poor penetration through thick scattering mediums (e.g. skin), which would limit the translation of this approach for continuous biomonitoring.

**Figure 3.**
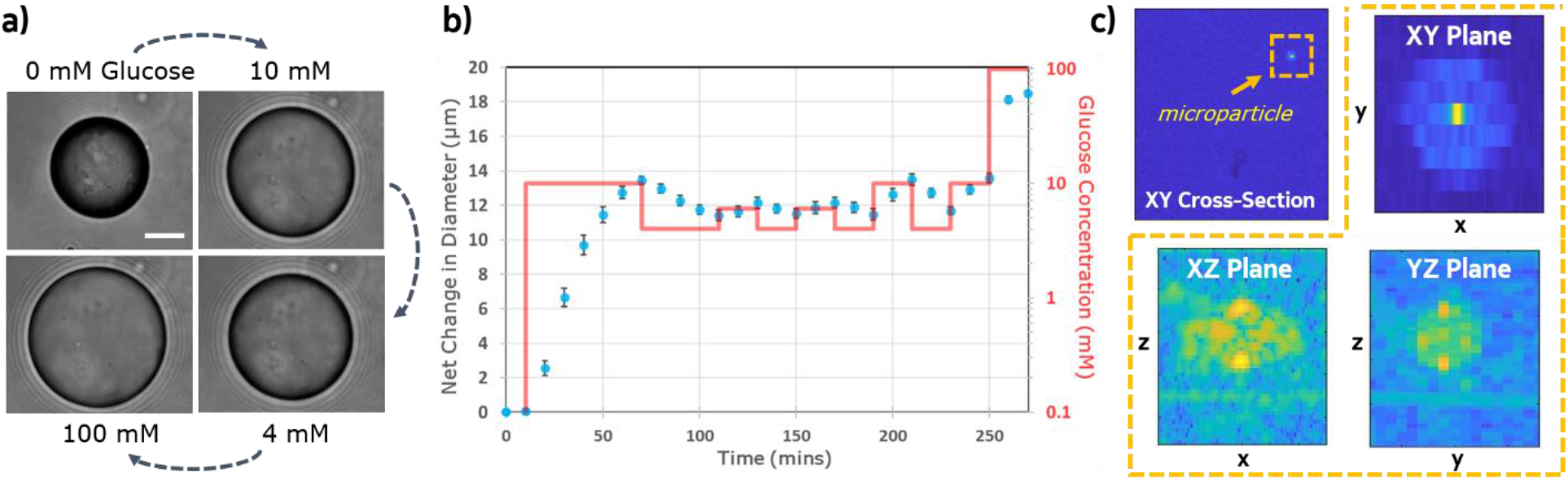
Spatiotemporal tracking of microparticle biosensors embedded in tissue-mimics exposed to constant glucose flow. (a) Microscopy images of microparticle swelling behavior to varying glucose concentrations. Scale bar: 20μm. (b) Plot of the net change in microparticle diameter size over time of microparticles embedded in a tissue-mimic exposed to a constant flow of varying glucose concentration *(red line).* The microparticle diameter was calculated by manual segmentation from images acquired using optical microscopy. The average microparticle starting size was about 55μm (n = 4). (c) OCT images displaying cross-sectional views of indicated microparticle *(dotted box)* in three different planes.

### Design and implementation of OCT for spatiotemporal biosensing

In contrast to optical microscopy, swept-source OCT (SS-OCT) is a well-established method for volumetric imaging in highly scattering mediums, with penetration depths greater than 1mm readily achievable in skin (see **Supplementary Note 1**). The method is based on low-coherence interferometry, in which a laser light beam from a tunable frequency swept laser source is directed to a sample and the backscattered light is compared to a reference beam to acquire a single depth scan (**Supplementary Fig. 6**). Single depth scans (i.e. A-scans) can be acquired at rates exceeding 100kHz, allowing for volumetric imaging of roughly 3mm x 3mm x 3mm volumes per second with 10μm resolution in all dimensions. Recent work to miniaturize these systems^33,34^ offers the potential of a portable, low-cost device for continuous *in vivo* monitoring, making SS-OCT an ideal candidate to acquire a non-invasive readout of microparticle biosensors.

OCT images of the flow cell were acquired over a lateral area of 1mm x 1mm using a custom fiber-based SS-OCT system incorporating a commercial swept-source laser centered at 1300nm. Microparticles displayed sufficient contrast to be identified in the acquired OCT images, and can be easily visualized in any spatial plane of interest (**Fig. 3c**). The axial resolution of our system was measured as 7μm (5.3μm in water), consistent with an optical bandwidth of 130nm.

Since size changes of the microparticles in physiologically-relevant concentrations of glucose were less than 10μm, which is less than the resolution of the OCT system, we developed two approaches to measure sub-resolution changes in microparticle diameter (see **Supplementary Note 2** for more details on both methods). The first method utilized Lorentzian peak fitting (**Supplementary Fig. 7**). Once each microparticle was identified, the A-scan with the strongest amplitude was selected. Since the microparticles were much larger than the OCT axial resolution, the top and bottom of the microparticle are clearly distinguishable in the A-scan. Each peak was fit to a Lorentzian line shape, from which the location of the peak position was calculated. The diameter of the microparticle is then just the difference in calculated peak position between the top and bottom surface of the microparticle.

The second method utilized spectroscopic fitting, which involved extracting the backscattered spectrum from raw OCT data using short-time Fourier transforms. The optical spectrum backscattered from a microparticle depends on the particle size, shape, and refractive index. Since the shape and refractive index of the microparticles are known *a priori*, the size can be determined by comparing experimentally measured backscatter spectrum to theory. To do this, we modeled the spherical microparticle as a Fabry-Perot optical cavity, wherein the incident laser light is confined to reflect between the top and bottom surface of the microparticle (**Supplementary Fig. 8**). The theoretical spectra calculated from the Fabry-Perot model were then autocorrelated to the experimentally-acquired spectrum to infer the microparticle size (**Supplementary Fig. 9 & Supplementary Fig. 10**). This method enabled small changes in microparticle diameter caused by glucose binding to be accurately inferred by monitoring its spectral backscatter (**Fig. 4a**). This method was benchmarked both on commercially-available polystyrene microparticles of known standardized sizes (see **Supplementary Note 3 & Supplementary Fig. 11**) as well as on multiple different sizes of glucose-responsive microparticles (**Fig. 4b & Supplementary Fig. 12**), with an accuracy of 1.2μm.

**Figure 4.**
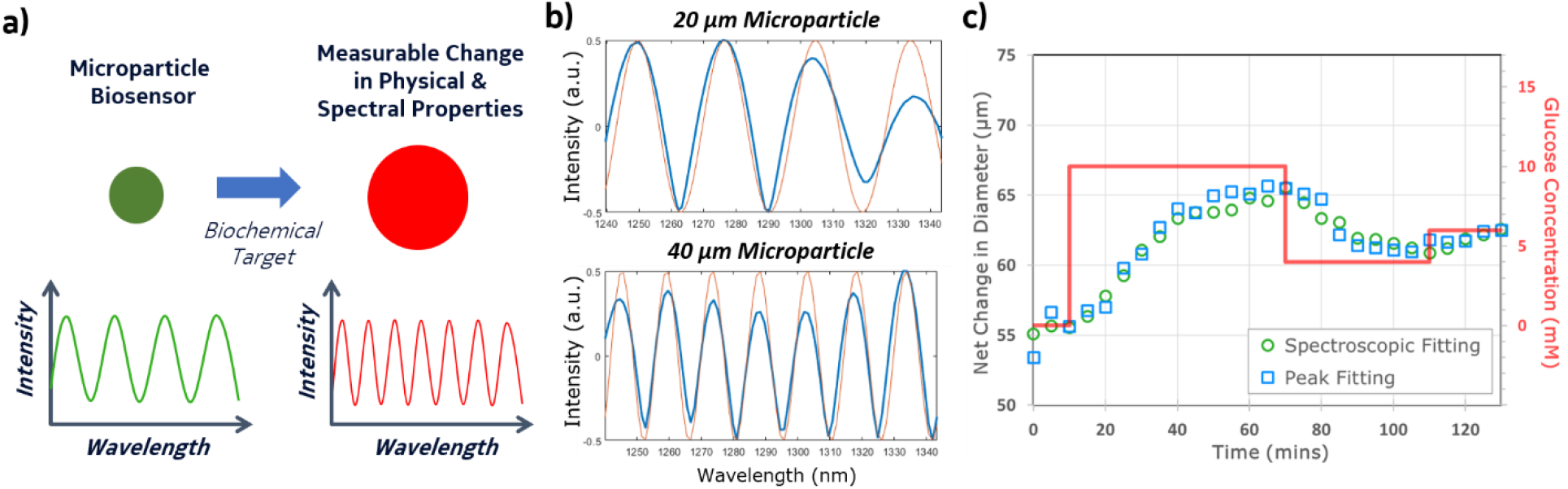
Monitoring microparticle biosensor response using spectroscopic OCT. (a) Biochemical target binding induces a physical change in the microparticle, resulting in a measurable optical change. (b) Overlay of experimentally acquired spectra (*blue*) and the predicted spectra (*red*) by autocorrelation with a Fabry-Perot optical cavity model, of two different sized microparticles embedded in a tissue-mimic. (c) Plot depicting the calculated size of the glucose-responsive microparticles using spectroscopic fitting (*blue circles*) and peak fitting (*red squares*), in response to varied glucose concentrations (*red line*) over time under constant flow conditions.

The flow cell setup described earlier was then used to monitor the microparticle response to glucose under continuous flow conditions using OCT. An initial concentration of 0mM glucose was applied for 10min, followed by a step increase to 10mM for 60min, a step decrease to 4mM for 30min, and then finally a step increase to 6mM for the final 15min. A region of interest (ROI) containing a single microparticle sensor was hand-segmented from the 4D OCT data, and analyzed using the peak fitting and spectroscopic method to determine its size at every time point. The two methods yielded nearly identical size estimates, as shown in **Fig. 4c**. With the step change in flow from 0mM to 10mM, the 55μm microparticle took approximately 35min to reach 90% of its final diameter. The microparticle then shrank in size when the flow was decreased to 4mM, and again increased in size upon the final step increase to 6mM. The estimated sizes closely follow the variations in glucose concentration within the flow cell, demonstrating reversable swelling and deswelling of the particle to easily distinguish physiologically-relevant concentrations for normal and hyperglycemic levels. As an early proof of concept, these are remarkable outcomes, which highlight the potential of our methodology to wirelessly track physiochemical changes of 3D-embeddable micron-scale sensors in response to biochemical fluctuations.

### Deep learning for automated tracking of microparticle biosensors

The methodology described so far requires significant human input in order to identify ROIs containing a microparticle within the 3D OCT time series data, which is a tedious and time-consuming process. The lack of automation in this step is one key barrier for ultimately achieving real-time, continuous biochemical monitoring using our approach. We addressed this shortcoming by implementing a deep learning approach based on convolutional neural networks (CNNs). The application of CNNs on large image datasets^35^ for object recognition is one of the most important breakthroughs for artificial intelligence research in the last decade^36^. CNN and its many variants have become especially useful for medical image analysis^37,38^. In particular, they have been extensively applied to analyzing magnetic resonance imaging (MRI) and computerized tomography (CT) images, with the goal of either classifying the presence/absence of disease state or segmenting specific areas of interest (e.g. tumors, organs)^39^. Although there are a few recent reports for employing CNNs on OCT images, the focus has been primarily in optometry for classifying retinal disease and segmenting tissue regions in the eye^40,41^. Moreover, there is limited work on applying CNNs to track microparticles^42,43^, especially in 3D volumetric data like those generated by OCT.

To completely automate our end-to-end pipeline for OCT-based biochemical monitoring, we designed a microparticle detection algorithm based on a 3D CNN architecture (**Fig. 5a**). The architecture follows the typical design used in computer vision^35,36,44^. We utilized convolutional layers with small filters of size 3×3×3 voxels, followed by rectified linear units (ReLU) and maxpooling layers of stride 2 to reduce the input dimensions (**Supplementary Fig. 13**). Since 3D CNNs have more parameters than the typical 2D CNNs used in image classification, we intentionally kept the filters small to reduce the number of parameters and prevent overfitting. We repeated this construct four times, followed by two linear layers (with ReLU activation in between) to reduce the output dimension to two with softmax activation for classification. The total number of trainable parameters in the CNN is 13,346. More details about the network architecture and the training are provided in the Methods section.

**Figure 5.**
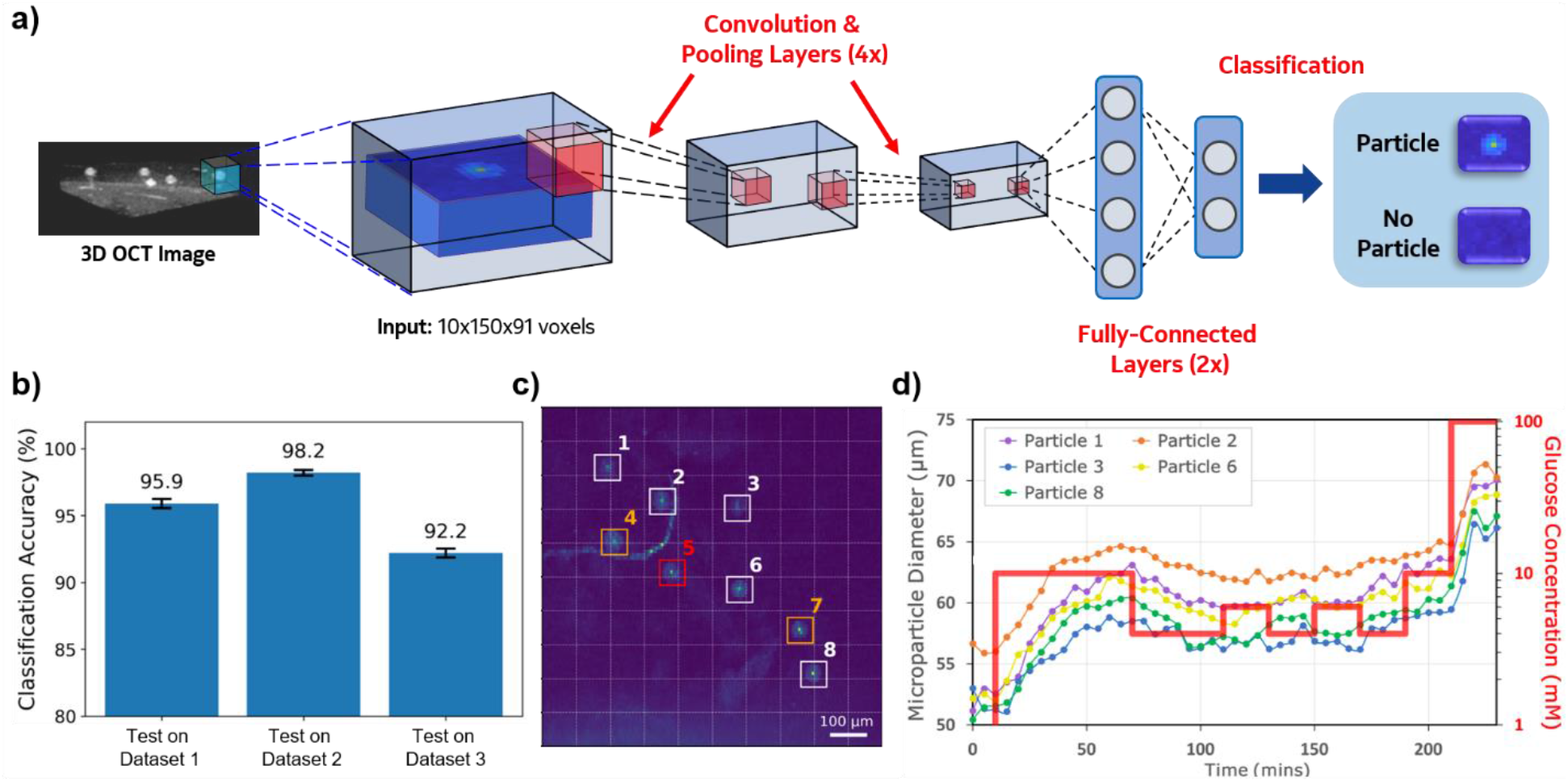
Automated microparticle tracking using deep learning. (a) 3D convolutional neural network architecture for microparticle classification, consisting of four convolutional layers followed by two full-connected linear layers. (b) Plot depicting the microparticle classification accuracy (presence/absence) across the different datasets. (c) Cross-sectional OCT image displaying microparticles that were successfully identified (*white & orange*) and missed (*red*) during classification. Microparticles indicated in *orange* displayed large fluctuations in size estimates which were physically improbable and were thus excluded from further monitoring. (d) Multi-particle monitoring of changes in microparticle size in response to varying glucose concentrations (*red line*) over time under constant flow conditions.

The 3D CNN was first trained to classify whether an input volume contains a microparticle or not. For this, the raw OCT volumetric image (1mm x 1mm x 4mm) was divided into sub-volumes of 100μm x 100μm x 594μm (10 x 150 x 91 voxels), whereby each sub-volume served as a distinct input into the network. The z-dimension of the sub-volume input was selected to contain the full depth range in which the microparticles were known to be confined (within the flow cell described earlier). Moreover, a given sub-volume was assumed to have at most one microparticle, which is a reasonable assumption considering the predefined experimental conditions (i.e. known size range of the microparticle batch, and the dilution used to achieve sparse spatial distribution within the tissue-mimic).

The training and testing data were acquired from three different sets of continuous glucose flow cell experiments (Dataset 1, Dataset 2 and Dataset 3). The volumetric scans in each experiment contained at least eight glucose-responsive microparticles and were captured every two to five minutes over a span of about three to four hours. We also included volumetric scans of samples containing microparticles with added background scatter (750-nm polystyrene beads) as training data to improve the robustness of the 3D CNN towards noise. We manually located the microparticles, and generated positive training examples by placing random bounding boxes around the microparticles. These random bounding boxes were created by shifting a bounding box (10×100 pixels) from the microparticle center by up to three pixels in the y-direction and up to forty pixels in the x-direction. This equated to a shift in about one-third the size of the bounding box in the corresponding dimensions (z-dimension was kept constant, as described above). A maximum of three such random bounding boxes were created for each microparticle. An equal number of negative training examples were generated by sampling bounding boxes from the background. Dataset 1, Dataset 2 and Dataset 3 contained 1504/1504, 2016/2016 and 3276/3276 positive/negative examples, respectively. The noisy OCT scan with added background scatter contained 368 examples. The CNNs were then trained using stochastic gradient descent.

We performed an independent evaluation of the 3D CNN classification accuracy for detecting the presence/absence of microparticles. To assess the classification accuracy, we tested on three independent datasets (Dataset 1, Dataset 2 and Dataset 3). When we tested Dataset 1 (1504 positive and 1504 negative), we trained the 3D CNN using Dataset 2 and Dataset 3, as well as the noisy OCT scan with added background scatter, and calculated the classification accuracy of the trained model on Dataset 1. We accordingly repeated this process for Dataset 2 and 3. In this way, we evaluated how well the 3D CNN transfers to microparticles in an unseen experiment. The 3D CNN was found to have an accuracy of about 95% in classifying 3D regions for the presence of a microparticle (**Fig. 5b & Supplementary Fig. 14**). We also considered 2D CNNs to detect microparticles in the x-y plane by working directly with a 2D input, wherein the intensities of the OCT scan were summed over the z-axis in the given sub-volume. However, this was not as accurate as the 3D CNN due to loss of information from the z-axis.

After confirming a high classification accuracy for identifying microparticles within the 3D sample, we then performed an end-to-end evaluation of the entire biosensing pipeline. It is worth noting that we have thus far built a highly accurate ‘classifier’ to determine the presence/absence of microparticles in a chosen input sub-volume using a 3D CNN. But to apply it to ‘locate’ microparticles in a given volume for monitoring sub-micron size changes of the identified microparticle, we needed a strategy for generating as well as assessing sub-volumes of the full OCT scan. There are many ways to divide the OCT scan into a grid of sub-volumes. In general, a microparticle located at the center of a sub-volume was found to be more easily identified with a 3D CNN compared to a microparticle located at the boundary. In turn, the choice of grid layout would affect the overall accuracy of the microparticle detection pipeline. We first divided the 1mm x 1mm cross-sectional area into a 10×10 grid (white dotted grid lines in **Fig. 5c**) for the 3D CNN to classify. The z-dimension of each sub-volume in the grid was kept constant (at 91 pixels, as described earlier), since it contains the entire A-scan of the microparticle required for size estimation. This helped to reduce the labeling effort in trying to find the depth of the microparticle. To handle microparticles lying at the boundaries of grid cells, we also use the 3D CNN to classify a 9×9 grid shifted by half the size of a grid cell in the x and y dimensions, and take the union on the set of microparticles identified with the original set. By taking the union of two sets of microparticles identified by separate classification on two different grid layouts, we can improve the detection of microparticles lying at the boundaries in either of the grid layouts. Although this can potentially lead to more false positives (non-microparticles), this problem was circumvented by employing a stringent filter on which microparticles were selected for further tracking based on expected size changes. In particular, we deemed any detected microparticle that changed in size by more than 10% in one time step (i.e. five mins) to be unreliable and eliminated them from the list of microparticles to be tracked, since such a response was found to be physically infeasible during the microparticle characterization studies described earlier. Such detected microparticles could be either false positives (a non-microparticle like background scatter noise), or true positives (actual microparticles) with a noisy A-scan such that we cannot estimate their sizes reliably.

As depicted in **Fig. 5c** for one of the selected datasets, seven of the eight microparticles were successfully identified by our 3D CNN (indicated as white and orange bounding boxes). The evolution of microparticle size (estimated using the peak fitting method as described in **Supplementary Note 2**) over time was plotted for all identified microparticles, wherein the microparticles indicated by white bounding boxes displayed the expected swelling and deswelling response to glucose as described above (**Fig. 5d**). Two of the microparticles (indicated by orange bounding boxes in **Fig. 5c**) were excluded since the change in size per time step was physically improbable (**Supplementary Fig. 15**). We prefer tracking a set of fewer but more stable microparticles because we only need to reliably estimate the sizes of a handful of microparticles to monitor the fluctuations in biochemical concentration.

This approach of identifying microparticles using 3D CNN is highly flexible and can be adapted through training with appropriate data to different types of microparticles in different tissue environments to monitor diverse biochemical targets. It is also fast and can be made to run in real-time on embedded systems with AI chip accelerations for different portable medical devices. The main difficulty in applying this approach is the collection of labeled training data, which can be expensive and time consuming. But it could potentially be circumvented through the use of synthetic data (e.g. microparticle data generated through a physical model such as Fabry-Perot) and through the sharing of labeled data in public repositories by different groups, as done in many other domains of applications for deep learning^35,45^.

## CONCLUSIONS

Overall, we have demonstrated a novel OCT-based biosensing approach, using glucose-responsive microparticles as a proof of concept. Since inception, OCT has been primarily used as a tool for structural imaging—initially for optometry and recently for dermatology. By combining the unique attributes of OCT imaging with biochemical-responsive polymeric microparticles and machine learning, we report one of the first demonstrations of OCT as a robust biosensing modality.

Our approach offers exceptional attributes for continuous biochemical monitoring applications. First, the microparticle biosensors are freely dispersible in any liquid medium, scaffold or even biological tissue. This provides the potential to monitor biochemicals in their native biological environments, without requiring complex wiring or tethering to the surface. In turn, the microparticle biosensors can be potentially deployed into the skin and thereafter imaged with OCT up to 1-2 mm inwards in a non-contact mode (**Supplementary Fig. 16**). Second, the ability to simultaneously acquire spatial and temporal information about the distributed biosensors allows for the response to be mapped in 3D over time, thus capturing the biochemical heterogeneity of a sample or tissue. Third, by decoupling the biorecognition component (i.e. microparticle) from the signal reporter (i.e. non-invasive and non-contact OCT imaging), our modular design imparts the flexibility to independently tune each sensing element. In turn, a versatile library of responsive microparticles can be envisioned that exhibit physiochemical changes based on the interaction of the microparticle-tethered biorecognition element (e.g. DNA/RNA, antibodies, aptamers, small molecules) with a target biochemical-of-interest^46-48^.

The combination of such a biosensor library with a reliable means of sensing readout like OCT enables our methodology to be tailored for a variety of prospective applications in continuous *in vivo* biomonitoring. For instance, an unmet need in chronic wound care is a facile means to acquire a biochemical readout of the healing progression, with minimal to no physical perturbation of the wounded tissue after the initial intervention^49^. We envision our methodology to be suitable for such an application, wherein biocompatible micron-scale sensors that sense the local changes in the wound biochemistry can be freely distributed within the wound, followed by remote non-contact monitoring in 3D over time using OCT. Moreover, recent efforts to miniaturize the OCT system from benchtop to chip-scale^33^ further empowers our approach, with broader implications to improve future healthcare monitoring in the clinic, at the home and on-the-go.

## METHODS

### Materials

All chemicals were of analytical grade and used without further purification. Acrylamide (AM) (99.5%), polyethylene glycol diacrylate (PEGDA) (MW: 575), paraffin oil, hexane (anhydrous, 95%), dextrose (D-(+)-Glucose), agarose, dimethyl sulfoxide (DMSO) (sterile-filtered, 99.7%) and monodisperse polystyrene microparticles (4.0 μm, 6.0 μm, 10.0 μm & 20.0 μm) were purchased from Sigma-Aldrich. 2-hydroxy-2-methylpropiophenone (2-HMP) (96%) and Span80 was purchased from TCI America. 3-(acrylamido)phenylboronic acid (APBA) was purchased from Boron Molecular. Sodium phosphate buffer (0.2 M, pH 8.5) was purchased from Alfa Aesar. Clear silicone sealant was purchased from Loctite. Porcine skin (non-sterile, 1.524 mm) was purchased from Stellen Medical.

### Equipment

A UV-spot curing probe (BlueWave QX4, 365 nm) was purchased from Dymax. A hand-held optical power meter (1830-R) was purchased from Newport. Syringe pumps (Legato 200 dual syringe infuse only) were purchased from KD Scientific. Chambered coverglass (Lak-Tek® II, #1.5 borosilicate, 8 wells) was purchased from Electron Microscopy Sciences. Hypodermic needles were purchased from Becton Dickinson. A low-profile, rubber platform vortex mixer was purchased from Thermo Scientific. The optical microscope (A1R Laser Scanning Confocal Microscope) was purchased from Nikon.

### Formulation of the precursor monomer solution

The precursor solution was prepared by first dissolving AM (62-72mol%) and PEGDA (3mol%) in deionized water. The phenylboronic acid derivative APBA (25-35mol%) was then dissolved in DMSO and added dropwise to the AM-PEGDA solution, followed by the addition of the 2-HMP photoinitiator (5wt%). The precursor solution was then mixed and passed through a 0.45 μm syringe filter. The total weight of AM, PEGDA and APBA was adjusted with respect to the volume to acquire ~18% w/v gels.

### Fabrication of microparticles using the vortex method

To form microparticle emulsions using the vortex method, an oil mixture consisting of paraffin oil and 1% Span80 surfactant was first prepared. About 1mL of this oil mixture was then transferred to a 11-mL borosilicate glass vial, followed by the addition of 50μL of the precursor solution. The aqueous precursor solution was transferred into the oil mixture with a pipettor, forming a single droplet immersed in oil at the bottom of the vial. The sample was then vortexed (1500-2500 rpm) for 40 seconds to allow for emulsion formation, followed by irradiation with UV light (365 nm) for 3 min. The microparticles were then transferred to Eppendorf tubes to purify by sequential washing (hexane, followed by phosphate buffer solution) and isolation with centrifugation (1000g, 2 min).

### Fabrication of microparticles using microfluidics

A three-inlet, one-outlet geometry microfluidic device was used to generate monodisperse hydrogel emulsions via the flow-focusing technique. The middle inlet was 20μm in width by 20μm in depth orifice sandwiched by two side inlets that were 100μm in width and 20μm in depth. During droplet generation, the precursor solution was infused through the middle inlet at flow rates between 0.2-1 μL/min. Simultaneously, sheath fluid (consisting of paraffin oil and 5wt% Span80 surfactant) was introduced through the two side inlets at flow rates between 20-40 μL/min to hydrodynamically pinch the precursor fluid at the orifice, dispersing the uniform droplets at desired sizes. Droplets with sizes ranging from 20 to 60 μm were generated with by adjusting the flow rates in described ranges. The droplets were then transported via the device outlet (500μm in length x 250μm in width x 20μm in depth) and connected tubing (30-gauge) to a transparent Eppendorf tube collector. For photo-polymerization, a UV-Spot curing probe was positioned 1 cm away from the outlet tubing exit, and a 5-mm diameter circular UV spot with an exposure intensity of 5 W/cm^2^ was continuously delivered to the exiting droplets to form spherical hydrogel microparticles. The microparticles were then purified by sequential washing (hexane, followed by phosphate buffer solution) and isolated with centrifugation (1000g, 2 min).

### Characterization of microparticle response to glucose

Glucose solutions of varying concentrations were generated in PBS (pH 8.5) by serial dilution, starting with a 100 mM stock. To assess the microparticle response to glucose under static conditions, about 10μL of the microparticle solution was transferred to the bottom of the well in chambered coverglass (Lak-Tek® II, 8-well). After waiting about 1 min for the microparticles to settle, the coverglass was placed on a holder in the optical microscope. After transferring 200μL of the desired glucose solution into the well, images were acquired every 30 s. To assess the microparticle response to glucose under constant flow conditions, a custom single-inlet and single-outlet flow cell was made. Briefly, the flow cell consisted of a PDMS slab with a through-hole recess measuring 25mm in length, 5mm in width, and 5mm in depth. The PDMS slab was irreversibly bonded to a glass slide forming an open flow cell. Prior to glucose solution infusion, an equal part mixture of 1% molten agarose (maintained at 75°C) and the microparticle suspension solution was mixed for five seconds, the resultant 0.5% mixture solution was then dispensed as a thin layer in the PDMS flow cell. The flow cell was immediately placed at 4°C for three minutes in order to harden the agarose gel matrix, and then filled with buffer solution. A clear silicone sealant was subsequently applied to the open edges of the flow cell, allowing the placement of a cover glass to form an air-tight closure for continuous flow from inlet to outlet. All experiments were conducted at room temperature.

### OCT imaging setup

OCT images of the flow cell were acquired over a lateral area of 1mm x 1mm using a custom fiber-based swept-source OCT system incorporating a commercial swept-source laser (Axsun; 1310nm center wavelength, 130nm sweep bandwidth, 100 kHz repetition rate). A galvo scanner was used for beam scanning along with a telecentric lens that provided 10um lateral resolution at the focal plane. The lateral dimension was raster scanned, with the 1mm x-direction significantly spatially oversampled for a total of 1500 A-scans per B-scan. The y-direction was scanned at 10 μm increments for a total of 100 pixels over 1 mm. A full scan took approximately 1.6 seconds and was repeated every two to five minutes. More details for the OCT setup are provided in **Supplementary Note 1**.

### Deep learning architecture

We employed a 3D CNN architecture as shown in **Supplementary Fig. 13**. It consists of multiple blocks of convolution and maxpooling layers, followed by linear layers that map to the classification output. It also follows the design principle of VGGnet^44^ that uses small filters and double the number of channels whenever there is a downsampling. The output of a block xk+1 can be computed from the output of the previous block xk as:

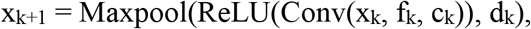

where Conv(x, f, c) is the convolution operator with a filter size f and c output channels, Maxpool(x, d) is max-pooling operator with d downsampling factors in different dimensions, and ReLU is the rectified linear unit. We use f_k_ = (3,3,3), (3,3,3), (1,3,3), (1,3,3), c_k_ = 8, 16, 32, 64, and d_k_ = (2,2,2), (2,2,2), (1,2,2), (1,2,2). There are two linear layers that reduce the dimension to 64 with ReLU and then 2 with softmax for classification (presence/absence of microparticle). We use the cross-entropy loss to train the network for classification. The 3D CNN is trained 40 epochs (40 passes through the training data) by stochastic gradient descent using a step size of 0.1 and a batch size of 128. The neural network was implemented using Tensorflow^50^. It required about four hours to finish the training using one NVIDIA P100 GPU.

## ACKNOWLEDGMENTS

We would like to thank Dr. Sanjay Patel, Dr. Marcus Weldon, Dr. Chris White, Dr. John D. Kim and Dr. William (Sean) Kennedy from Nokia Bell Labs for inspiring conversations about this work and potential applications. We would also like to thank Dr. Michael Crouch from Nokia Bell Labs for his valuable inputs and feedback on the manuscript.

## AUTHOR CONTRIBUTIONS

S.S. conceived the project and performed the experiments. M.S.E. designed and implemented the optical coherence tomography methods. M.Z. conducted the microfluidic-based synthesis of microparticles. C.Y. and H.K. designed and implemented the deep learning algorithms. S.S., M.S.E. and C.Y. wrote the final manuscript.

