## Supplementary Information for "Microparticle-based Biochemical Sensing Using Optical Coherence Tomography and Deep Learning"

[a] Nokia Bell Labs, New Providence, NJ, U.S.A.

[b] Current address: Department of Computer Science and Engineering,
New York University, New York, NY, USA

***Correspondence:**

**Dr. Shreyas Shah**


### TABLE OF CONTENTS


| |
|---|
| **Supplementary Fig. 1.** Schematic of reversible covalent binding to a phenylboronic acid derivative |
| **Supplementary Fig. 2.** Images depicting microparticles synthesized using the vortexing method |
| **Supplementary Fig. 3.** Scale and setup of the microfluidics devices for microparticle generation |
| **Supplementary Fig. 4.** Plot depicting the net response of different sized-microparticles |
| **Supplementary Fig. 5.** Experimental setup for flow cell experiments |
| **Supplementary Note 1: Basic Principles of Swept-Source OCT**<br>➢ **Supplementary Fig. 6.** Schematic drawing of the swept-source OCT setup |
| **Supplementary Note 2: Microparticle Size Estimation Methods**<br>➢ **Supplementary Fig. 7.** Overview for size estimation using the Peak Fitting method<br>➢ **Supplementary Fig. 8.** Schematic of modeling microparticles as Fabry-Perot optical cavities<br>➢ **Supplementary Fig. 9.** Schematic of acquiring depth-resolved spectroscopic information<br>➢ **Supplementary Fig. 10.** Overview for size estimation using the Spectroscopic Fitting method |
| **Supplementary Note 3: Benchmarking Microparticle Size Estimation Methods**<br>➢ **Supplementary Fig. 11.** Validation of size estimation using commercial microparticles<br>➢ **Supplementary Fig. 12.** Experimental and predicted spectra of different-sized microparticles |
| **Supplementary Fig. 13.** 3D convolutional neural network architecture |
| **Supplementary Fig. 14.** Confusion matrix for the 3D CNN classifier |
| **Supplementary Fig. 15.** Glucose response of microparticles excluded by the automated pipeline |
| **Supplementary Fig. 16.** Cross-sectional OCT image of microparticles in pig skin |
| **Supplementary Table 1.** Percentage change in microparticle size and the corresponding time constant |

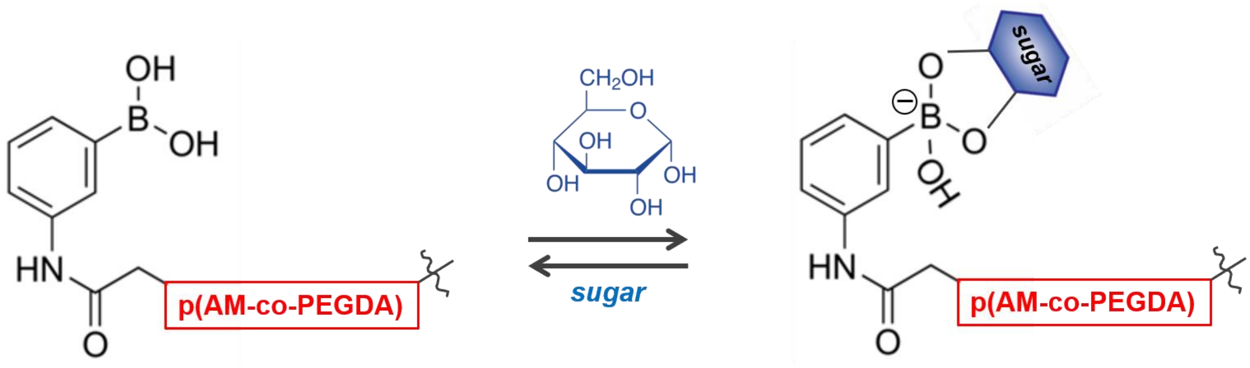


**Supplementary Figure 1.** A representation of the reversible covalent binding of the anionic boronate species from the phenylboronic acid derivative with the *cis* diol group from glucose molecules. The glucose-binding 3-(acrylamido)phenylboronic acid is copolymerized with a poly(acrylamide-*co*-poly(ethylene glycol) diacrylate) p(AM-*co*-PEGDA), forming a hydrogel matrix.

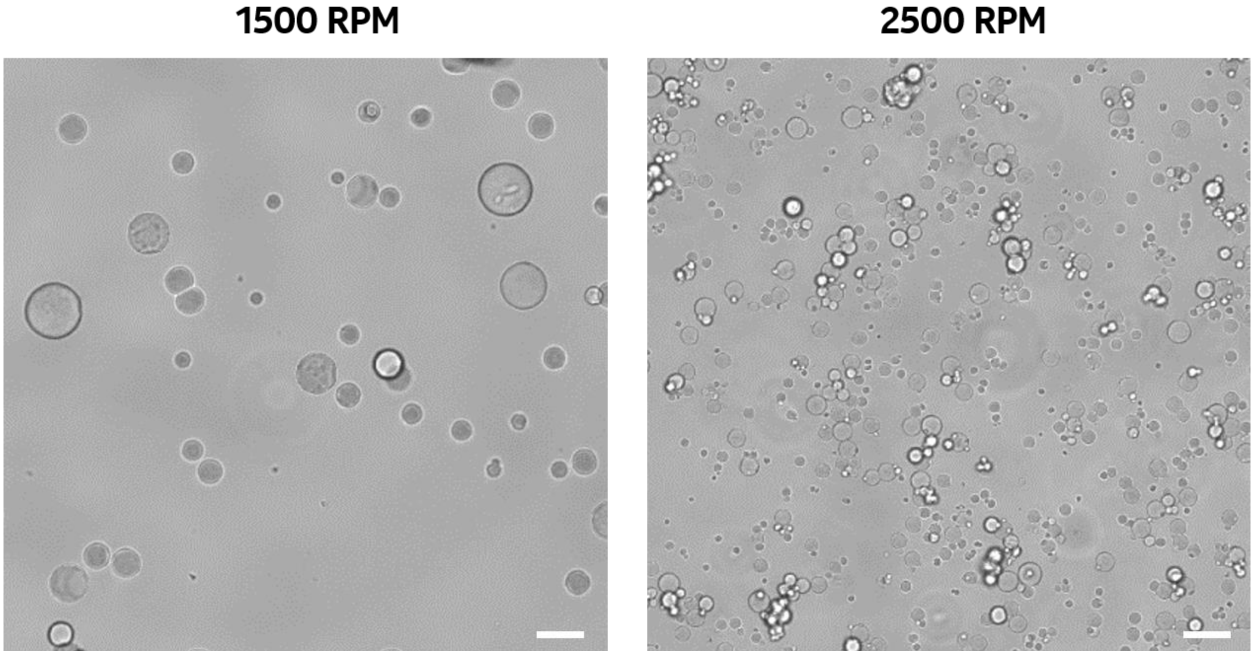


**Supplementary Figure 2.** Images depicting glucose-responsive microparticles synthesized using the vortexing method at two different speeds (1500 RPM and 2500 RPM). Microparticles with varying sizes can be generated at a given vortexing speed, wherein higher speeds result in smaller sizes. Scale bars: 25 μm.

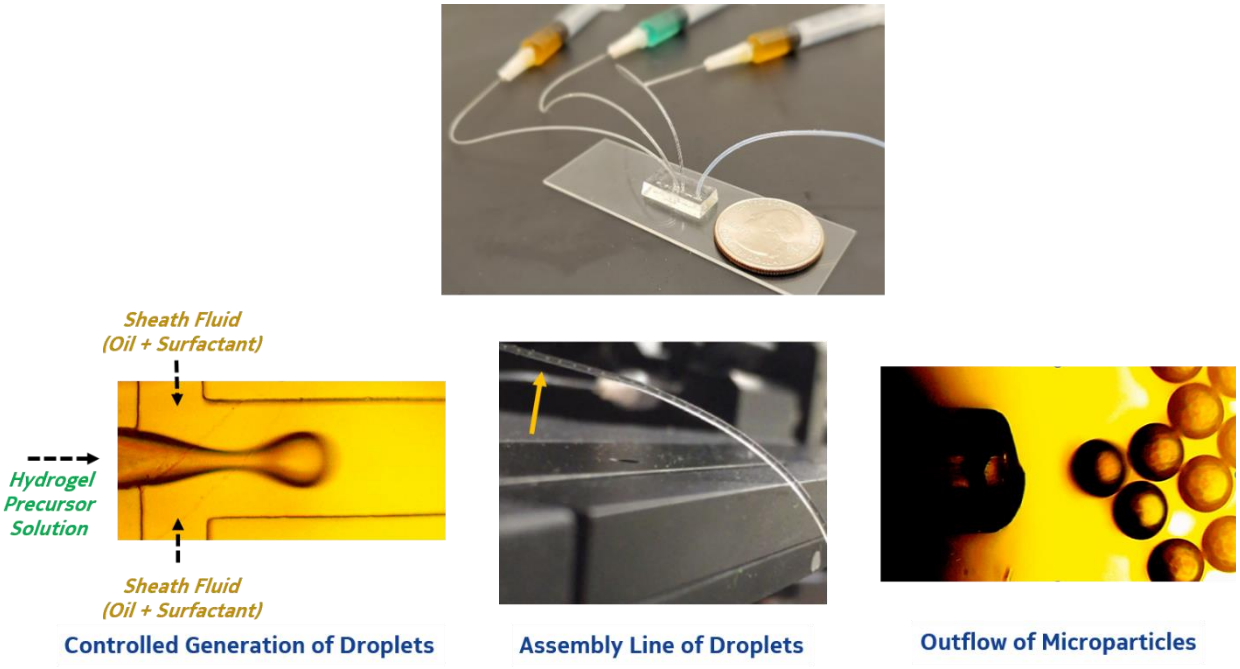


**Supplementary Figure 3.** Images depicting the scale and setup of the microfluidics devices for microparticle generation.

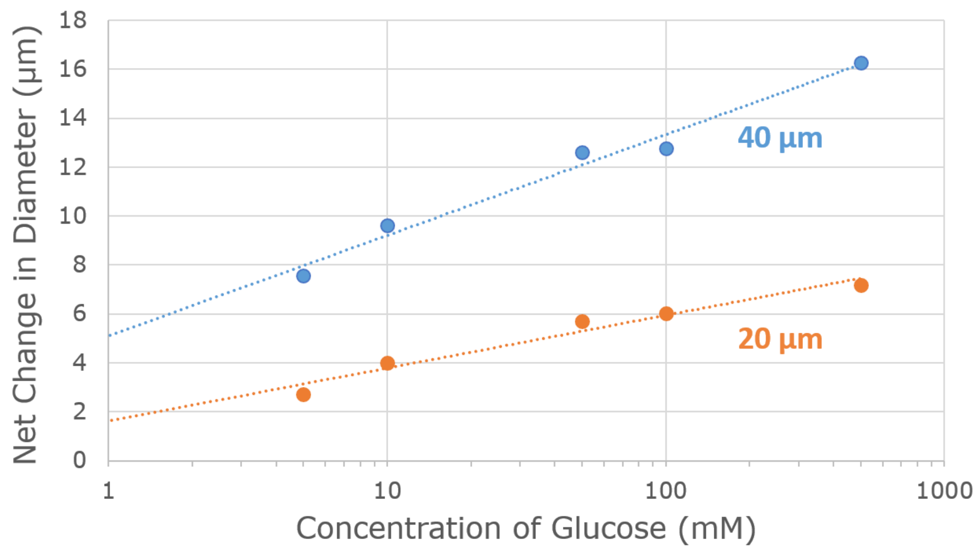


**Supplementary Figure 4.** Plot depicting the net response of different sized-microparticles (20µm and 40µm) synthesized using microfluidics, at varied concentration of glucose.

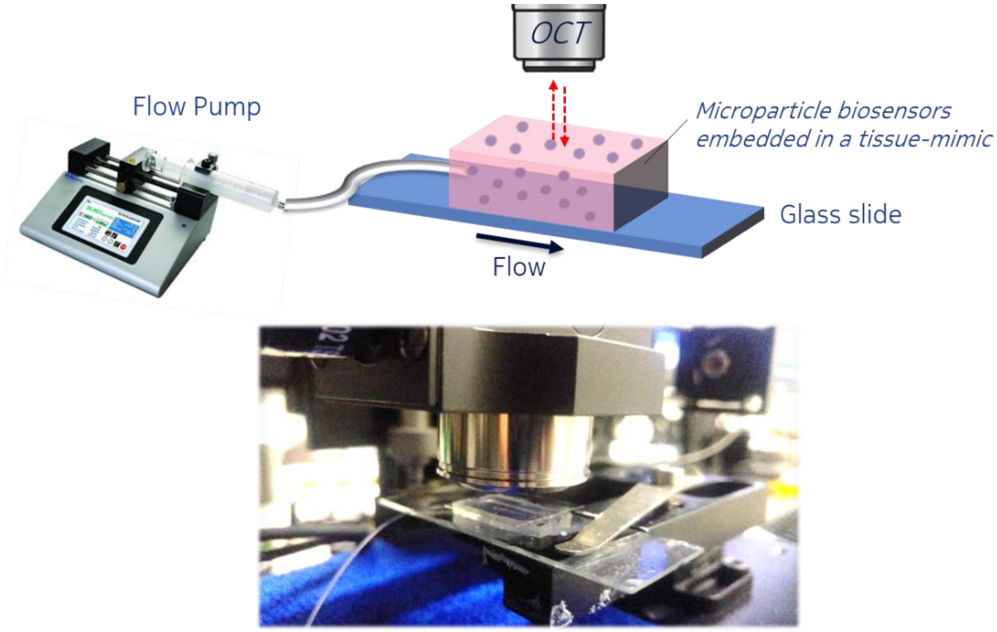


**Supplementary Figure 5.** Experimental setup for monitoring microparticle biosensors embedded in a tissue-mimic exposed to a constant flow (1 mL/min) of glucose solution. The microparticle embedded tissue-mimic was positioned in the flow cell, which was fixed in position below the OCT scan lens.

### **SUPPLEMENTARY NOTE 1:**
### BASIC PRINCIPLES OF SWEPT-SOURCE OPTICAL COHERENCE TOMOGRAPHY (SS-OCT)

Swept-source optical coherence tomography (SS-OCT) is an optical ranging technique that utilizes a self-homodyne interferometer design to detect path length changes between a sample and the reference arm (**Supplementary Fig. 6**). In our custom-built system, we utilize a commercial fiber-coupled swept-source laser (Axsun), which continuously changes its output optical frequency in time at a repetition rate of 100kHz. The total optical bandwidth scanned over is 130nm with a center frequency of 1300nm (i.e. approximately 23THz bandwidth centered at 230THz).

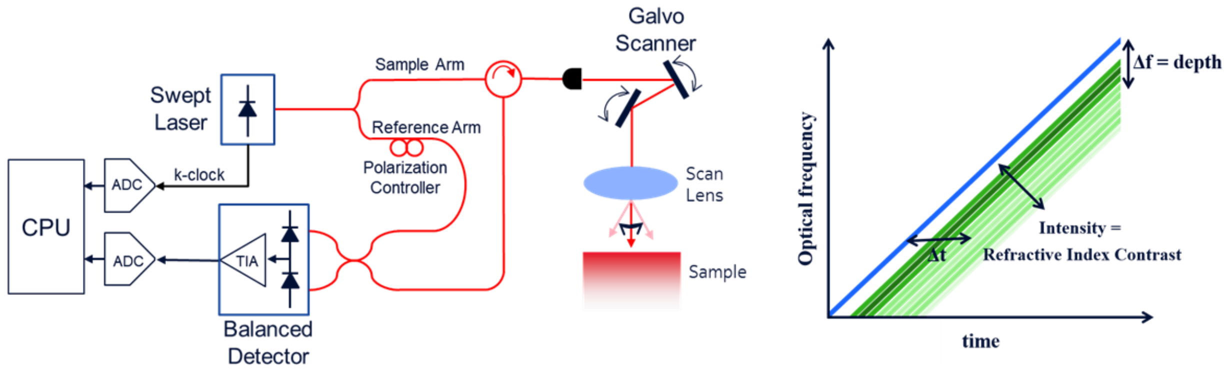


***Supplementary Figure 6.*** *Schematic drawing of the swept-source optical coherence tomography (OCT) setup (left) and the corresponding schematic of optical self-homodyne (right). The laser light beam from a tunable frequency swept laser source is split: one beam goes into free-space and is reflected back from the target (target arm) and the second beam stays local and beats against the reflected beam. The difference in frequency between the two beams is proportional to the distance traveled (depth) and the intensity is indicative of the refractive index contrast in the target.*

The system uses a Mach-Zehnder interferometer architecture, where the laser is split into two paths using a 90/10 fiber splitter. One path contains a delay arm of known length and the other path is sent to the sample being measured. The sample path contains a galvo scanning mirror (Thorlabs) that allows the laser to be raster scanned over two dimensions, followed by a scan lens (Thorlabs LSM02) which enables telecentric scanning of the sample. The system achieves approximately 20µm lateral resolution in focal plane.

The backscattered light from the sample is collected with the same scan lens, coupled back into fiber and mixed with the reference arm on a balanced photodiode. A circulator is used to isolate forward and backwards propagating light in the sample arm and to avoid excess losses a simple splitter would incur.

The output of the balanced diode is digitized using a 1GSa/s 16bit ADC. A fixed length interferometer built into the Axsun laser provides an electronic k-clock signal, which is also digitized. The swept-source laser produces a non-linear scan which must be corrected before image reconstruction. This is done by taking a Hilbert transform of the digitized k-clock signal and using the unwrapped optical phase to resample the OCT signal to linear-phase. Dispersion mismatch between the reference and sample arm is compensated for digitally by multiplying the resampled OCT signal by a fixed phase rotation (calibrated using a mirror in the sample path). An FFT of the OCT signal is then taken to produce the final OCT intensity versus depth plot.

---------------------------------------------------------------------------------

### **SUPPLEMENTARY NOTE 2:**
#### MICROPARTICLE SIZE ESTIMATION METHODS

In this section, we describe in detail the two methods we used to estimate microparticle size. For both approaches, the first step was locating a region-of-interest (ROI) defined as a 3D bounding box containing exactly one microparticle from the full 3D OCT scan. Once an ROI was identified, the intensities are summed up across the z-axis (depth scan or A-scan) at each point in the x-y plane and the point of maximum intensity is used as the center location of the microparticle.

For the first method, we employed Lorentzian peak fitting (**Supplementary Fig. 7**). In the depth scan identified above as the center of the microparticle, the two largest peaks were found, which correspond to the top of the microparticle and the bottom of the microparticle. A Lorentzian lineshape was then fit to each peak using the intensity values of the two voxels towards the microparticle-side of the peak (i.e. inside the microparticle) and ten voxels on the external-side of the peak (i.e. outside the microparticle). From this fitting, the depth corresponding to the top-surface peak and bottom-surface peak was calculated. Because thirteen total points were used in fitting the lineshape, the resolution of the peak value is significantly better than the depth resolution itself. The microparticle optical diameter was then calculated as the top-surface peak minus the bottom-surface peak. Finally, the physical diameter was determined by dividing the calculated optical diameter by the refractive index of the microparticle, which was known *a priori*.

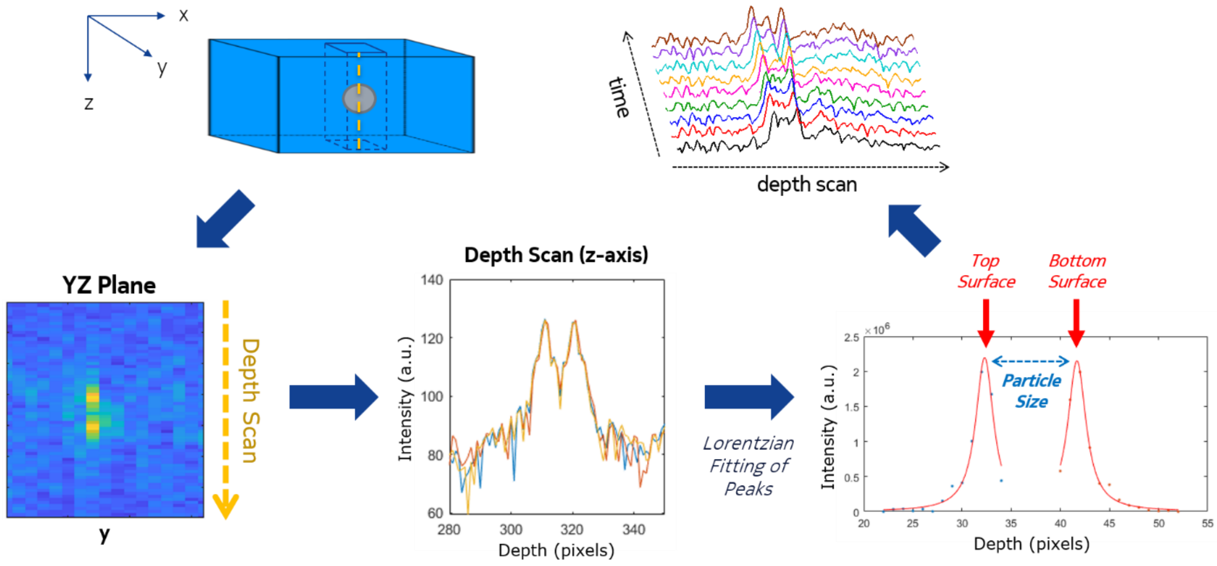


***Supplementary Figure 7.*** *Overview of the steps for estimating the microparticle size based on the Peak Fitting method. The depth scan (A-scan) corresponding to the highest intensity of the microparticle in the XY plane is plotted. The two prominent peaks in the depth scan profile, corresponding to the top and bottom surface of the microparticle, are fitted using a Lorentzian function and the difference is used to calculate the diameter of the microparticle. This allowed tracking changes in the microparticle size over time.*

For the second method, we employed a spectroscopic fitting approach. The light backscattered from a spherical particle can be modelled in several different ways. In our case we looked at two simple models: (a) Mie scattering, which gives the back reflected spectra within the acceptance cone of our scan optics for a given particle size and refractive index, and (b) a Fabry-Perot (FP) model (**Supplementary Fig. 8**), where the top and bottom surface of the microparticle are modeled as partial reflectors and the total back-reflected light can be analytically calculated based on the refractive index of the particle, the refractive index of the surrounding medium, and the particle diameter. The FP model is reasonable for OCT-based backscatter due to the very low numerical aperture of our scan lens, which results in a near-planewave excitation pattern.

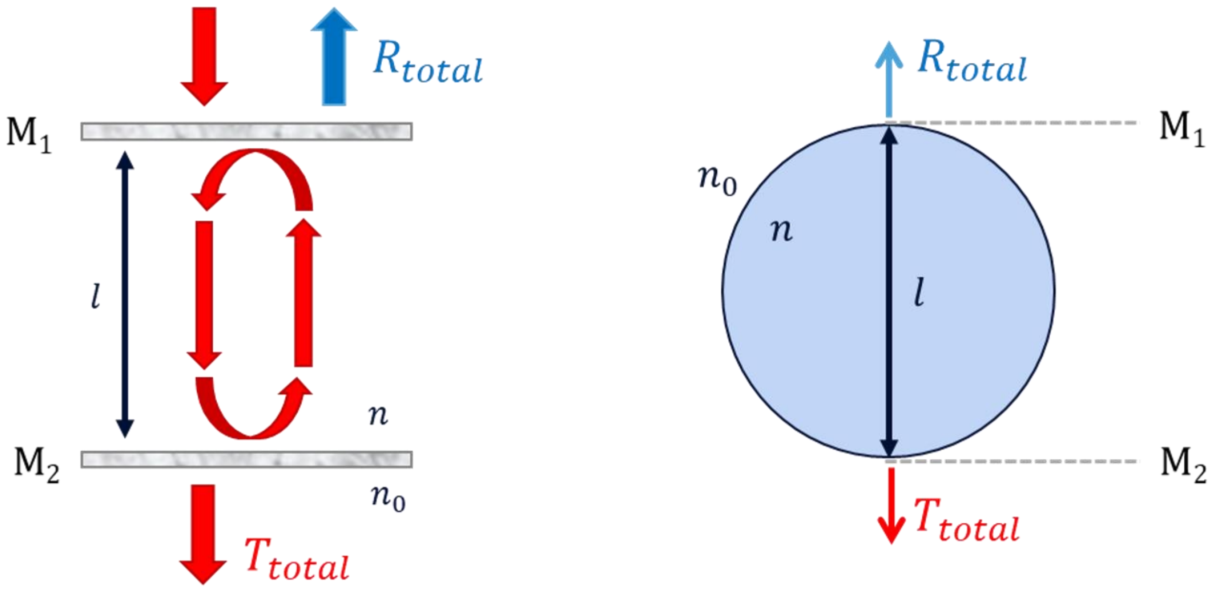


***Supplementary Figure 8.*** *Conventional Fabry-Perot optical cavity (left) consisting of two partially reflecting parallel mirrors, in which optical waves can pass through the cavity only when in resonance with it. The spherical microparticle (right) was modeled as a Fabry-Perot optical cavity, wherein the incident laser light is confined to reflect between the top and bottom surface of the microparticle.*

Both models were tested using commercially-available polystyrene microparticles of known refractive index and size. The FP model was found to be more accurate, especially for larger-sized microparticles (see **Supplementary Note 3**). For this reason, an FP model was used to estimate microparticle size of the glucose-responsive microparticles, which had an approximate initial size of ~50µm.

To calculate the backscattered spectrum, the brightest A-scan in the ROI was converted back into a time-domain signal using an inverse Fourier transform. A spectrogram was then calculated for this signal using a sliding FFT window of 100 points and a 95% overlap. The resultant spectrogram, as depicted in **Supplementary Fig. 9**, provides a plot of backscattered wavelength versus depth.

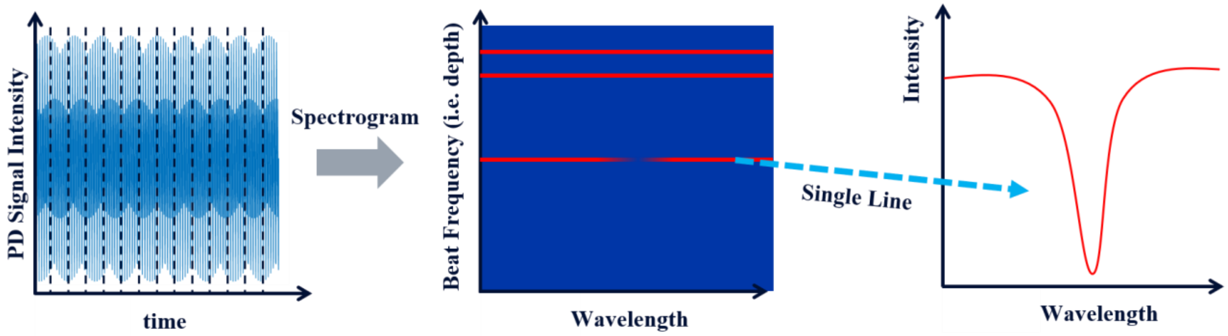


***Supplementary Figure 9.*** *Depth-resolved spectroscopic information is acquired by measuring the spectral content of backscattered light from the raw OCT signal. Short-time Fourier transforms of the OCT signal provides wavelength-dependent variations at a given depth (i.e. beat frequency), which can give material-specific information.*

Two spectral signals were then extracted from this 2D plot: the signal spectra (i.e. the backscattered waveform originating from the depth where the microparticle ROI is located) and the background spectra (in our case, the backscattered signal from a coverslip sitting on top of the sample). The normalized backscattered signal was then calculated as the signal spectra divided by the background spectra as shown in **Supplementary Fig. 10**.

The expected normalized backscatter spectrum was calculated for microparticles in the range of 10 to 80μm in 70-nm step sizes. These theoretical spectra were then autocorrelated with the measured normalized backscattered spectrum—the diameter corresponding with the theoretical spectra most highly correlated to the experimentally-measured spectra was assigned as the estimated microparticle size. A typical autocorrelation plot is shown in the bottom right panel of **Supplementary Fig. 10**.

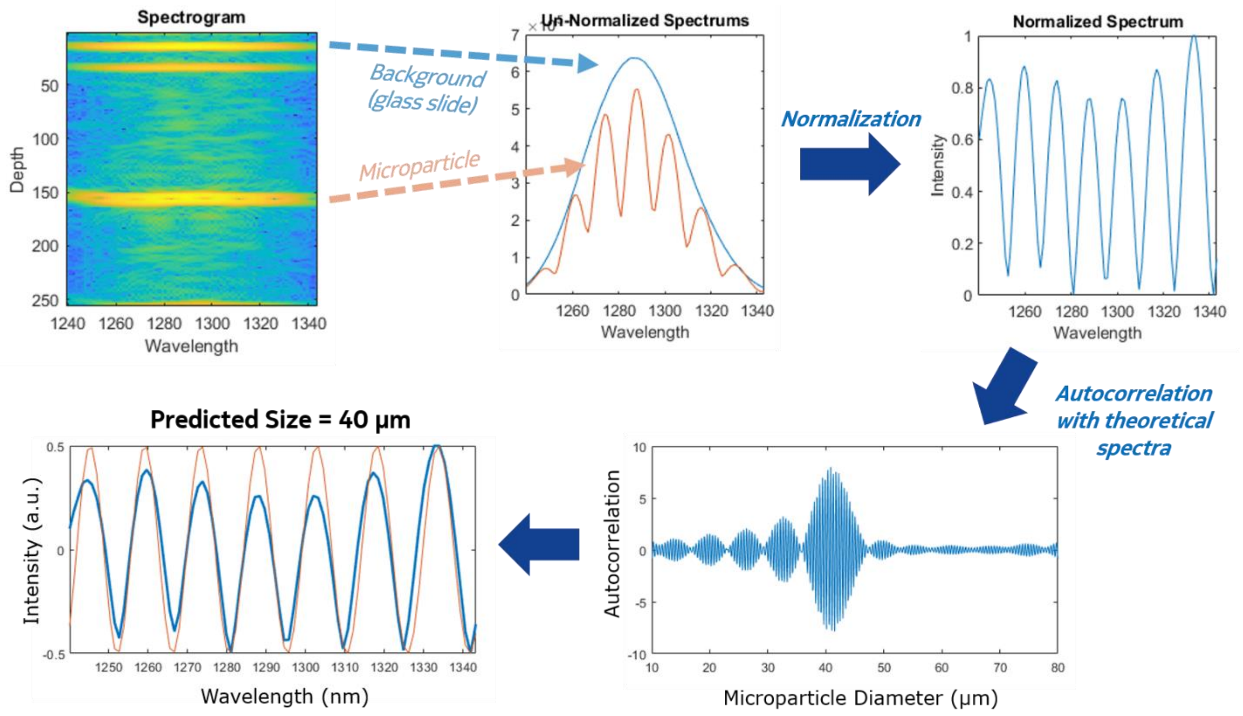


***Supplementary Figure 10.*** *Overview of steps for estimating the microparticle size based on the Spectroscopic Fitting method. A depth-resolved spectrogram is acquired by measuring the spectral content at the depth scan (A-scan) corresponding to the highest intensity of the microparticle in the XY plane. The spectrum corresponding to the microparticle (which shows an oscillation in the amplitude) is extracted from the spectrogram and normalized with the background glass spectrum. The theoretical spectra calculated from a Fabry-Perot model for a range of microparticle sizes is then autocorrelated to the experimentally acquired spectra to predict the microparticle size.*

### **SUPPLEMENTARY NOTE 3:**
### BENCHMARKING MICROPARTICLE SIZE ESTIMATION METHODS

To validate the spectroscopic fitting method (i.e. backscatter spectrum-fitting) for estimating microparticle size, experiments were conducted on commercial polystyrene microparticles (Sigma-Aldrich; refractive index = 1.56) of known size embedded in a PEG gel. Four different diameter microparticles were used: 4µm, 6µm, 10µm, and 20µm, each with a diameter error of less than 300nm. OCT images of these particles were captured, and their sizes were determined using the method descripted in **Supplementary Note 2**, using both a Mie scattering model and a FP model. The results, shown in **Supplementary Fig. 11**, showed that both Mie and FP models yield accurate results at small particle sizes, however at larger particle sizes (d > 10µm) the Mie scattering model significantly under-estimated particle size.

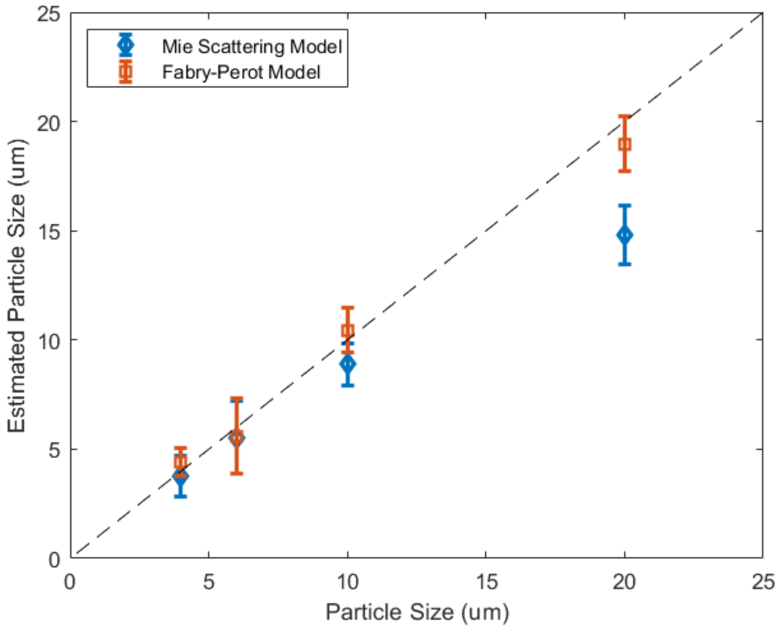


***Supplementary Figure 11.*** *Plot comparing the measured size (calculated from the Fabry-Perot model) versus the actual size of commercially available standardized polystyrene microparticles. The mean measured absolute size error is less than 1µm with a standard deviation less than 1µm.*

Over all the size ranges, the FP model gave a mean absolute diameter offset error of 0.6µm with a standard deviation of 1.2µm. This is significantly below the depth resolution of our system, which is measured as 7µm (in air). By comparing *en-face* optical microscopic measurements with OCT estimations, we found the FP model works well even for microparticles exceeding 20µm as shown in **Supplementary Fig. 12**.

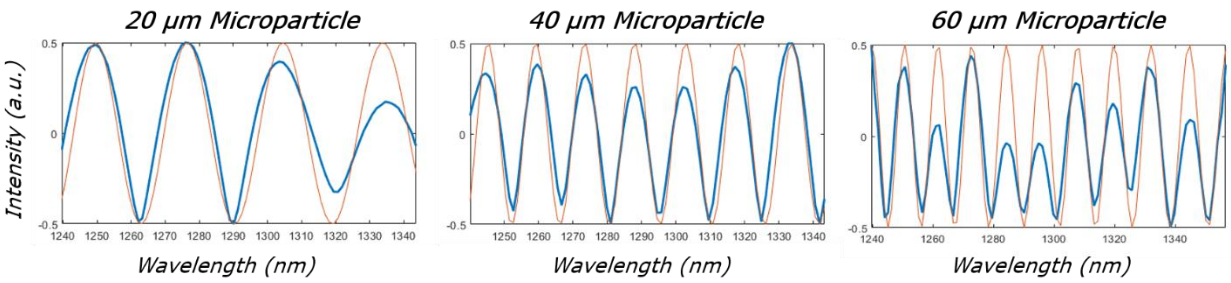


***Supplementary Figure 12.*** *Overlay of experimentally acquired spectra (blue) and the predicted spectra (red) by autocorrelation with a Fabry-Perot optical cavity model, of three different sized glucose-responsive microparticles embedded in a tissue-mimic.*

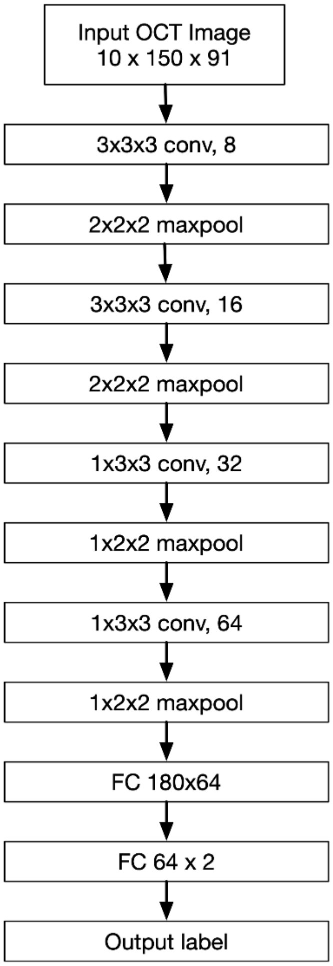


**Supplementary Figure 13.** Architecture diagram for the 3D convolutional neural network. There is a set of rectified linear units (ReLU) after each of the convolutional layers in the diagram. Conv, convolutional layer; FC, fully connected layer.

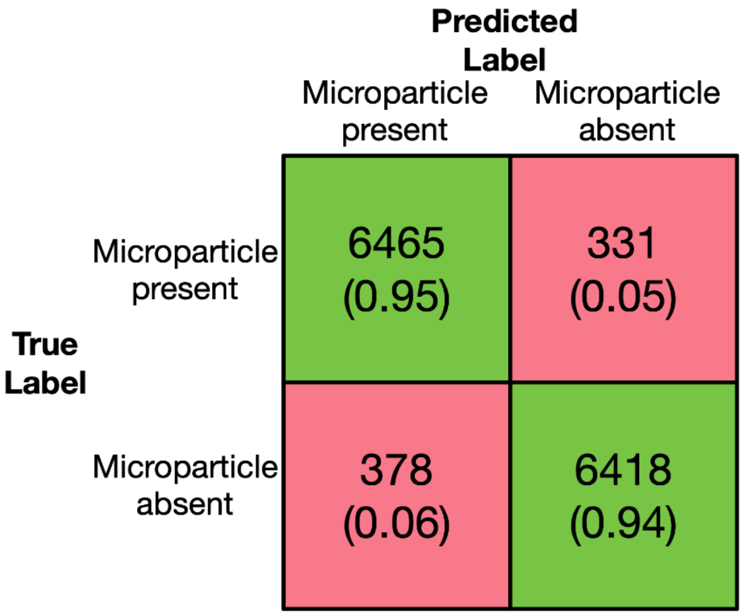


**Supplementary Figure 14.** Confusion matrix for the 3D CNN classifier on Test Set 1, 2 and 3 (combined). The numbers in parentheses (from left to right, top to bottom) are the true positive rate, false negative rate, false positive rate, and true negative rate, respectively. The overall accuracy is about 95%.

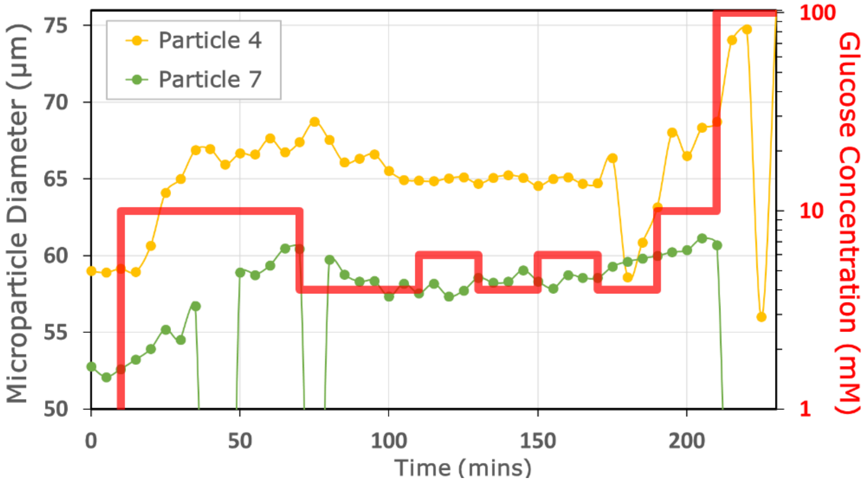


**Supplementary Figure 15.** Glucose response plot of microparticles that were excluded by the automated pipeline, due to large fluctuations in size estimates observed at multiple time steps which are physically improbable.

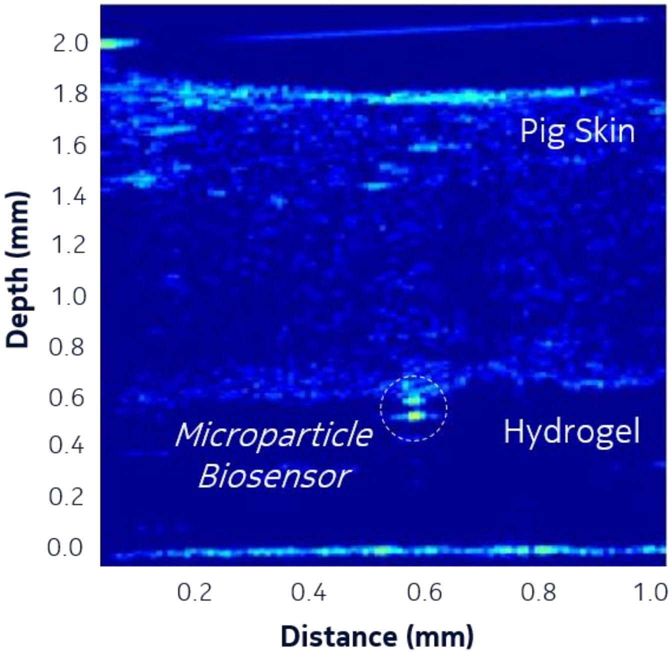


**Supplementary Figure 16.** Cross-sectional OCT image showing a microparticle biosensor embedded in a hydrogel tissue-mimic positioned below pig skin.

**Supplementary Table 1.** Table outlining the average percentage change in microparticle size and the corresponding time constant ($\tau$) for three different batches exposed to 100 mM glucose (1xPBS, pH 8.5) under static conditions. The molar ratio of the phenylboronic acid to acrylamide was varied, while keeping the poly(ethylene glycol) constant. The time constant, $\tau$, was calculated as the time taken for a microparticle to change from 10% to 90% of total size change.

| AM : PEGDA : APBA | Avg % Change in Size | $\tau$ (min) |
|---|---|---|
| 72 : 3 : 25 | 25.1 ± 3.0 | 3 |
| 67 : 3 : 30 | 32.3 ± 5.5 | 5 |
| 62 : 3 : 35 | 45.2 ± 4.5 | 8 |
